# The Use of Artificial Intelligence In Magnetic Resonance Imaging of Epilepsy: A Systematic Review and Meta-Analysis

**DOI:** 10.1101/2025.09.19.677393

**Authors:** Judy Chen, Ella Sahlas, Yigu Zhou, Natalie Chen, Jim Xie, Farhan Wadia, Sienna Armstrong, Lorenzo Caciagli, Alexander G. Weil, Roy W. Dudley, Dewi V. Schrader, Andrea Bernasconi, Neda Bernasconi, Boris C. Bernhardt

## Abstract

**Background:** The application of artificial intelligence (AI)/machine learning (ML) to MRI can be a powerful tool to streamline clinical decision-making, yet variability amongst MRI sequences and algorithms have hindered appropriate assessment of reliability and generalizability.

**Methods:** We conducted a systematic review and meta-analysis of the ability of current AI/ML models operating on MRI data for: 1) epilepsy diagnosis, 2) temporal lobe epilepsy lateralization, 3) lesion localization, and 4) post-surgical outcome prediction. Searches were conducted across PubMed, Medline, and Embase databases from inception until January 1, 2025. We selected studies that employed AI/ML models trained on any MRI modality to classify at least ten patients across the four main objectives for qualitative assessment, and further included in the meta-analysis if they reported an accuracy rate. The primary outcome of the meta-analysis was the overall accuracy of AI/ML models trained on MRI data. The secondary outcome was the concomitant risk of bias evaluation using PROBAST.

**Findings:** We identified 158 studies for qualitative evaluation and 127 studies for inclusion in the meta-analysis. AI/ML on multimodal MRI could accurately distinguish epilepsy patients from healthy controls (overall accuracy: 88% [85-90]), lateralize temporal lobe epilepsy (90% [87-93]), localize epileptogenic lesions (82% [74-88]), and predict post-surgical seizure-freedom (83% [78-87]). Overall, a high risk of bias remains in the literature; participant bias remained high across all outcomes (64-87%), as well as predictor (88-100%) and analysis (83-100%) bias. Outcome bias was low only for AI/ML studies predicting post-surgical outcomes.

**Interpretation:** Our results support promising accuracy of AI/ML models in epilepsy diagnostics and prognostics but remain highly susceptible to bias in participants, predictors, outcome, and analysis domains which limits current translation to routine clinical practice. We encourage closer interdisciplinary collaboration between clinical and scientific groups to improve validation studies based on thorough study design, analysis, and reporting.

## INTRODUCTION

Epilepsy is one of the most common neurological disorders, affecting >60 million people worldwide.^1^ In a third of people with epilepsy, seizure control cannot be achieved through pharmacotherapy. Considering the adverse effects of ongoing seizures and anti-seizure medication on health, cognition, and psychosocial functioning, current clinical care guidelines recommend referral of these patients to specialized epilepsy units for comprehensive pre-surgical investigation. Randomized trials and retrospective studies in adolescents and adults have shown that surgical treatment consistently leads to higher rates of seizure freedom than continued medical therapy.^2–4^ However, at present, only a small subgroup of potential surgical candidates receives surgery each year, even in high-income countries. An average of 20 years pass between the first seizure and epilepsy surgery for pharmacoresistant patients, a timespan associated with significant risks and burden for patients, families, and health care systems.^5–7^ Moreover, despite epilepsy surgery being the most effective treatment to achieve seizure control and restore quality of life for pharmacoresistant patients, it is not failproof. Indeed, 30-40% of patients who undergo surgery will experience seizures post-operatively.^8,9^ These challenges motivate the need to improve diagnostic and prognostic methods.

Magnetic resonance imaging (MRI) has revolutionized the evaluation and management of pharmacoresistant epilepsy, playing a pivotal role in the non-invasive detection and characterization of epileptogenic lesions.^10–13^ A lesion on MRI is the most reliable indicator of the surgical target and its complete surgical removal is a robust predictor for post-operative seizure freedom. Temporal lobe epilepsy (TLE) related to mesiotemporal alterations and epilepsy related to focal cortical dysplasia (FCD) remain the most prevalent diagnoses among patients with pharmaco-resistant focal epilepsy, and patients with these diagnoses benefit from advanced MRI imaging. Indeed, accurately lateralizing TLE and precisely localizing FCDs and other focal epilepsies are essential to successful post-surgical outcomes, yet both remain clinical challenging. In addition to limitations due to the lack of optimization of imaging protocols, these lesions may be subtle and can thus be overlooked by conventional visual exam^10,14^ Presently, artificial intelligence (AI) and machine learning (ML) have been increasingly leveraged to support key applications throughout the clinical workup: epilepsy diagnosis, TLE lateralization, lesion localization (including FCD localization), and prediction of post-surgical seizure freedom.

Current models remain mostly limited to few tertiary or quaternary care centers and have not been translated broadly. Moreover, heterogeneity in algorithms, MRI sequences, scanner types, and patient cohorts casts uncertainty on the design of an optimal model that can capture the complexity of the clinical challenges presented by the heterogeneity of epilepsy cases.^15–17^ For example, it remains unclear whether simpler ML models with few parameters perform similarly to more complex deep learning models, or whether AI/ML approaches perform similarly across different MRI sequences. The extent of heterogeneity across these factors and its influence on model performance has not been systematically investigated. Furthermore, the validity and generalizability of reported performance metrics have not been extensively evaluated. Given that an optimal, standardized approach to validating AI/ML models is required for improved and robust translation to routine clinical practice, a comprehensive review of current AI/ML-driven approaches in MRI for epilepsy is needed.

The objectives for this systematic review and meta-analysis were twofold. First, we assessed the performance of current AI/ML models to conduct major tasks: 1) diagnose epilepsy, 2) lateralize TLE, 3) localize the epileptic lesion, with a specific focus on FCD detection, and 4) predict post-surgical outcomes. Second, we evaluated the robustness of results and systematically appraised current literature to highlight sources of bias and to enable the identification of key barriers that prevent integration into routine clinical practice.

## METHODS

This review adhered to the Preferred Reporting Items for Systematic Reviews and Meta-analyses (PRISMA) guidelines and was registered on PROSPERO (CRD42023474405) prior to analysis.^18^

### Search strategy and study selection

We searched the Embase and PubMed databases from inception to January 1, 2025. Additional details of search terms and strategy are provided in **Figure 1**. The reference lists of included studies and relevant reviews were manually reviewed to include publications not captured from database searches. All titles/abstracts and full-text articles were independently screened by 2 reviewers (JC and SA or YZ). Disagreements were resolved through discussion and, if necessary, consultation with a third reviewer (JX). Publications using AI or ML classification methods on MRI-based images/features reporting on at least 1 of the 4 major clinical epilepsy applications listed were included: 1) diagnosis of epilepsy or differentiation between people with epilepsy and other populations, 2) TLE lateralization, 3) lesion localization, and 4) predicting post-surgical outcomes in pharmacoresistant patients. Studies that did not report any one of the evaluation metrics (*i*.*e*., sensitivity/specificity, positive predictive value/negative predictive value, F1 score, AUC under ROC, or raw accuracy rate), with less than 10 included patients, without full-text publications, or without original data were excluded.

**Figure 1.**
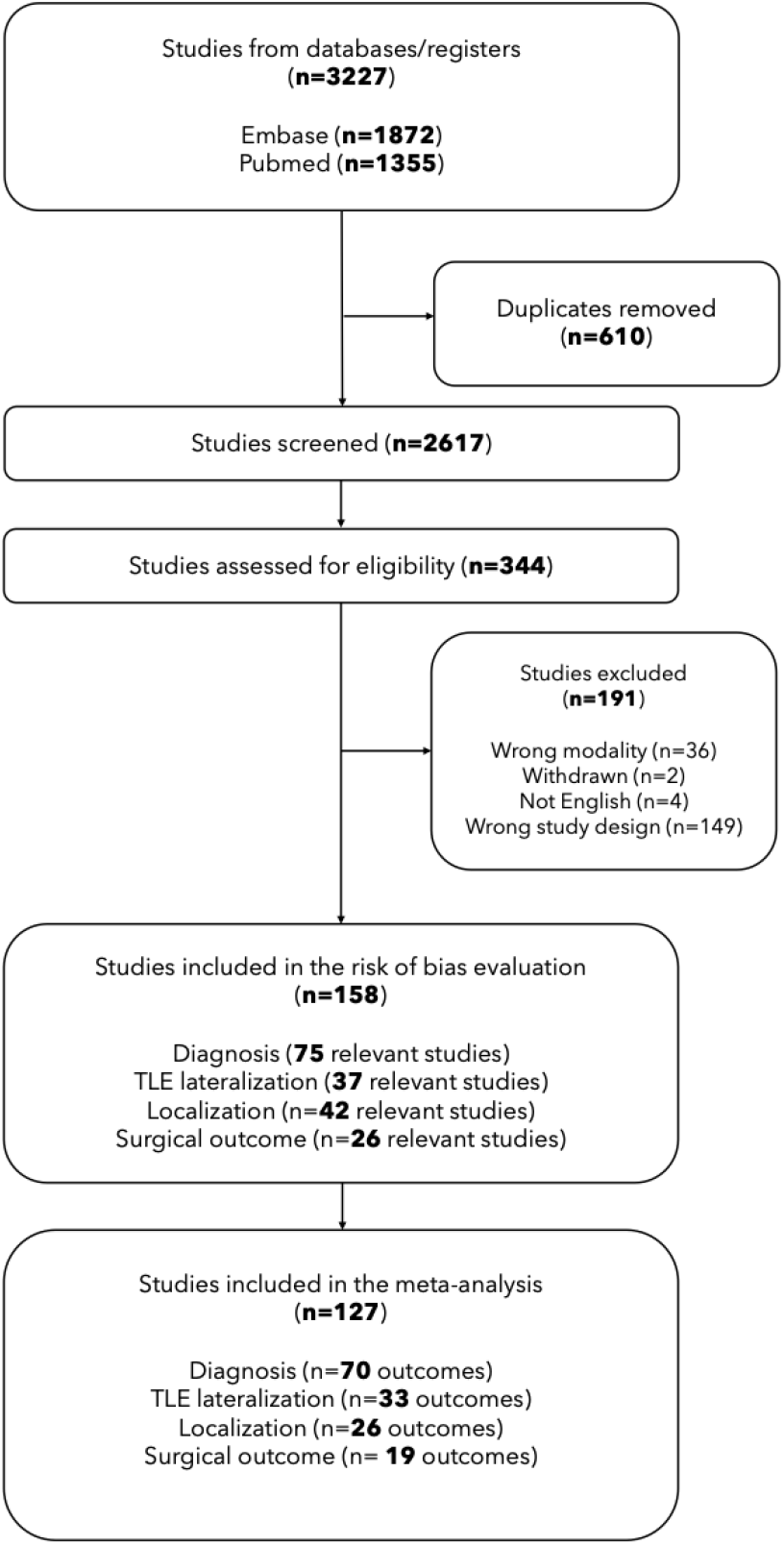
Preferred Reporting Items for Systematic Reviews and Meta-Analyses (PRISMA) flow diagram. A total of 3,227 studies were initially identified, with 158 studies ultimately included in the review after screening for duplicates, relevance, and eligibility and 127 in the meta-analysis. The included studies were categorized by topic (diagnosis, localization, lateralization, and surgical outcome) with the number of papers and corresponding number of outcomes reported for each topic. Note that some studies were relevant to multiple different topics and may also report multiple outcomes across differing populations or techniques.

### Data extraction and risk of bias assessment

Four reviewers (JC, FW, JX, NC) independently extracted data in duplicate. The following were extracted: (1) study characteristics (name of first and corresponding author, publication year, sample size), (2) demographic and baseline variables (patient and control cohorts’ age, sex, duration and/or age of onset of epilepsy, drug-resistance status, post-surgical seizure freedom, MRI-negative status), (3) data acquisition variables (MRI magnet strength, modality, data input), (4) ML characteristics (feature selection process if applicable, type of algorithm), and (5) evaluation metrics (sensitivity/specificity, positive predictive value/negative predictive value, F1 score, AUC under ROC, or raw accuracy rate). Two authors were assigned to each article; any discrepancy was resolved through discussion. The meta-analysis included only studies that reported an accuracy rate under their evaluation metrics. For studies that reported multiple applications with differing cohort groups (*e*.*g*., TLE lateralization of both MRI-negative and MRI-positive patients), all corresponding accuracy rates were included, while for studies that reported multiple accuracy rates across different algorithmic designs, only the one reporting the highest accuracy was recorded. This resulted in a higher number of reported accuracy rates than the number of studies. Risk of bias of studies were evaluated using the Prediction Model Risk Of Bias Assessment Tool (PROBAST), categorized across four domains (participants, predictors, outcome, and analysis) with an overall judgement of low, high, or unclear risk.^19,20^ Each quality assessment was conducted by at least two independent researchers (JC, ES, JX), and conflicts were resolved via discussion.

### Statistical analysis

Analyses were performed with the *metafor* package (version 4.4.0) in R (version 4.3.1, R Project for Statistical Computing).^21^ We examined the distribution of accuracy rates for all outcomes. Significance was defined as a 2-tailed *p*-value < 0.05.

## RESULTS

Of 3227 records retrieved, 158 studies were included in the qualitative and risk of bias analysis.^e1-e158^ Overall study characteristics are summarized in **Fig S1**. 127 studies were included in the quantitative meta-analysis (**Figure 1**).

### Identifying epilepsy

Of the 75 total studies, 71 attempted to classify epilepsy patients and healthy controls while 4 attempted to do so with diseased controls instead. There were 85 reported accuracy rates derived from 70 studies included in the meta-analysis. Overall, 11,410 participants (n=5,621 patients; M/F/NR 4112/4054/-, mean age: 28·5 years, mean reported disease duration: 15·4 years) yielded a mean accuracy rate of 88% (CI 85-89%, **Figure 2A**). The predominantly used ML technique was support vector machines (SVM; n=54), followed by neural networks (NN; n=13) and logistic regression (LR; n=7). Most studies used multimodal scans (n=26), with the most common unimodal scan being T1-weighted (T1w) MRI sequences (n=24) followed by diffusion MRI (n=19) and fMRI (n=17).

**Figure 2.**
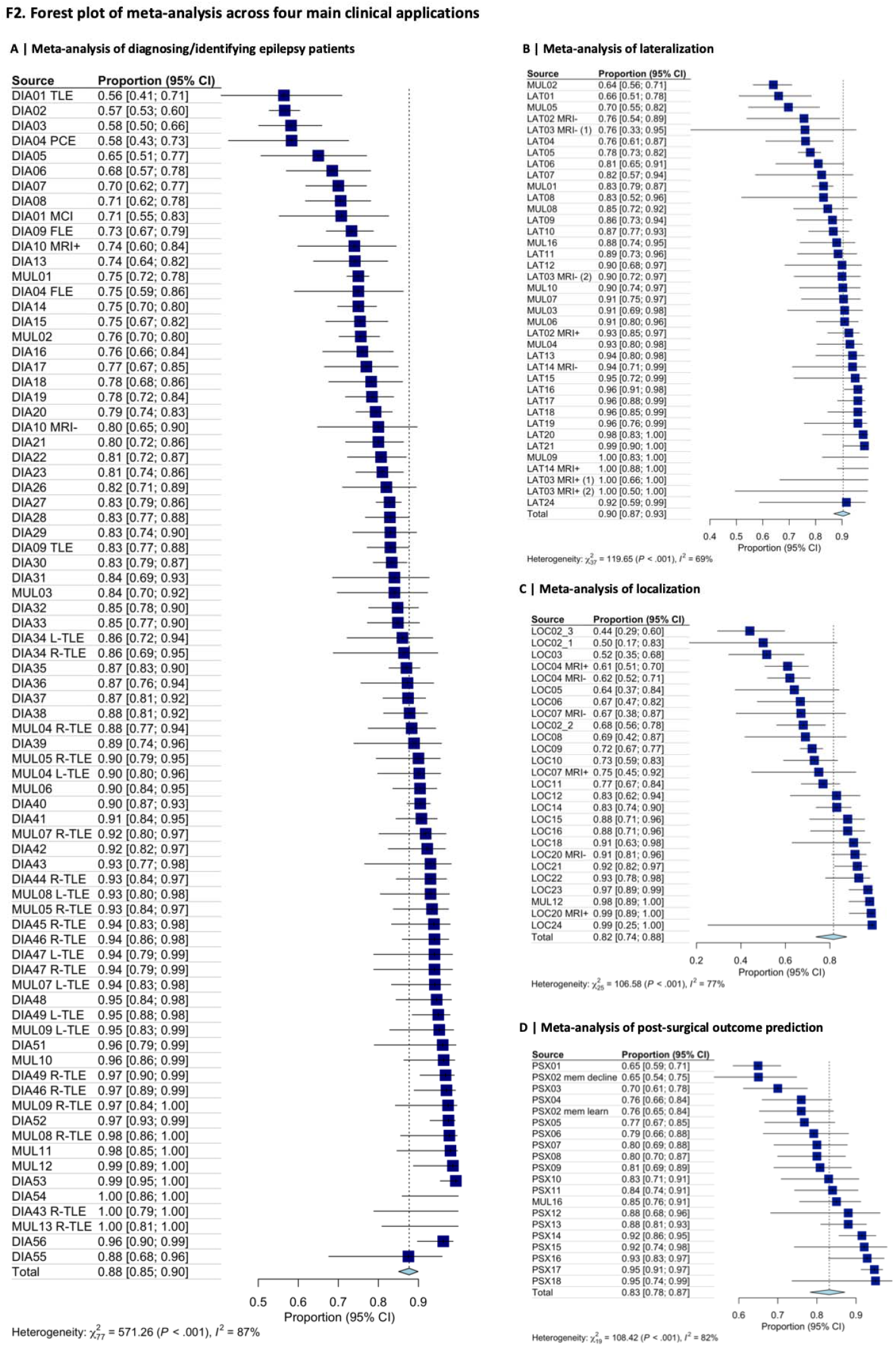
Forest plots of the meta-analyses for the four main outcomes. All recorded accuracy rates are listed and labelled according to the outcome (DIA = diagnosis, LAT = lateralization, LOC = localization, and PSX = post-surgical outcome). If a publication is part of multiple outcomes, MUL is used, and multiple cohorts are separated by left-(L)/right-(R) sided epilepsy, lesion class (malformations of cortical development [MCD]), or specific diagnosis (temporal lobe epilepsy [TLE], focal cortical dysplasia [FCD], MRI visible/positive [MRI+], etc.), which are also listed. The largest cohort size is displayed at the upper right and an icon representing the cohort size proportio against the maximum cohort size is shown across each listed accuracy rate. (A) For distinguishing epilepsy patients from other groups, the overall accuracy rate across 84 studies was 88% (CI 85-89%) across a total cohort size of 5275 epilepsy patients. (B) For lateralizing temporal lobe epilepsy, the overall accuracy rate across 37 studies was 90% (CI 87-93%) across 2303 patients. (C) For localizing the epileptic lesion, the overall accuracy rate across 2 studies was 82% (CI 73-91%) across 940 patients. (D) For post-surgical outcome prediction, the overall accurac rate across 20 studies was 83% (CI 78-87%) across 1410 patients.

### Lateralizing TLE

33/37 studies reported a total of 38 accuracy rates that were used in the meta-analysis. The mean accuracy rate was 90% (CI 87-93%, **Figure 2B**) across 2,396 total participants (n=1,071 patients; M/F/NR 1760/2198/-, mean age 33·8 years, mean disease duration: 18·0 years). SVM and discriminant analysis (DA) were the most common algorithm types used (n=20, 13). The most common MRI sequences used were T1w (n=16), multimodal combinations (n=10), and diffusion (n=8). Restricting the analysis to “MRI-negative” patients (n=555 patients; M/F/NR 199/230/-, mean age 35·5 years, mean disease duration: 17·3 years), the mean accuracy rate was 90% (CI 85-94%).

### Localizing focal epilepsy lesions

26/42 studies reported a total of 32 accuracy rates and were included in the meta-analysis; of these 42 studies, 25/42 studies included FCD patients exclusively. There was a total of 7453 participants (n=1742 patients; M/F/NR 2734/1214/-, mean age of patients: 21·9 years, mean disease duration: 9.76 years) included, and studies reported an overall accuracy of 82% (CI 73-91%, **Figure 2C**). Accuracy was evaluated at the subject level and did not incorporate false-positive extra-lesional findings, which may differ from the sensitivity and specificity of the corresponding models. Three studies limited inclusion to “MRI-negative” patients, while two included both MRI-negative and MRI-positive cases; all other studies (n=32) either did not report MRI-negative status or included MRI-positive scans only. Most studies employed a neural network framework (n=13) on multimodal MRI data, with the majority being a combination of T1w and FLAIR sequences (n=14).

### Predicting post-surgical outcomes

25 studies predicted seizure freedom (*i*.*e*., Engel class) after surgical intervention, while one study attempted to predict post-surgical memory learning and decline. Most studies (n=23) included TLE patients, while one included drug-resistant focal epilepsy patients and two included FCD patients only. There were 20 reported accuracy rates from 19 studies. The meta-analysis yielded an overall accuracy rate of 82% (CI 78-87%, **Figure 2D**) encompassing 1,820 total participants (n=1410 patients, M/F/NR 744/803/-, mean age: 32·7 years, mean reported disease duration: 17·3 years). The most common two algorithm types were SVM and DA (n=8, 4), and algorithms were most commonly trained on multimodal data containing T1w and diffusion MRI sequences (n=6).

We investigated whether reported accuracy rates for each outcome were associated with key study characteristics, namely cohort size, algorithm type, and MRI modality, using linear mixed effects models. Only larger cohort sizes were significantly associated with lower accuracy rates (**Figure 3A**), and this effect was most pronounced for diagnostic accuracy and post-surgical outcome prediction, exhibiting moderately negative correlations (r = -0·45, -0·51, p < 0.01). While associations to cohort size were also negative for localization and lateralization performance, these effect were not significant (r = -0·34, -0·2).

**Figure 3.**
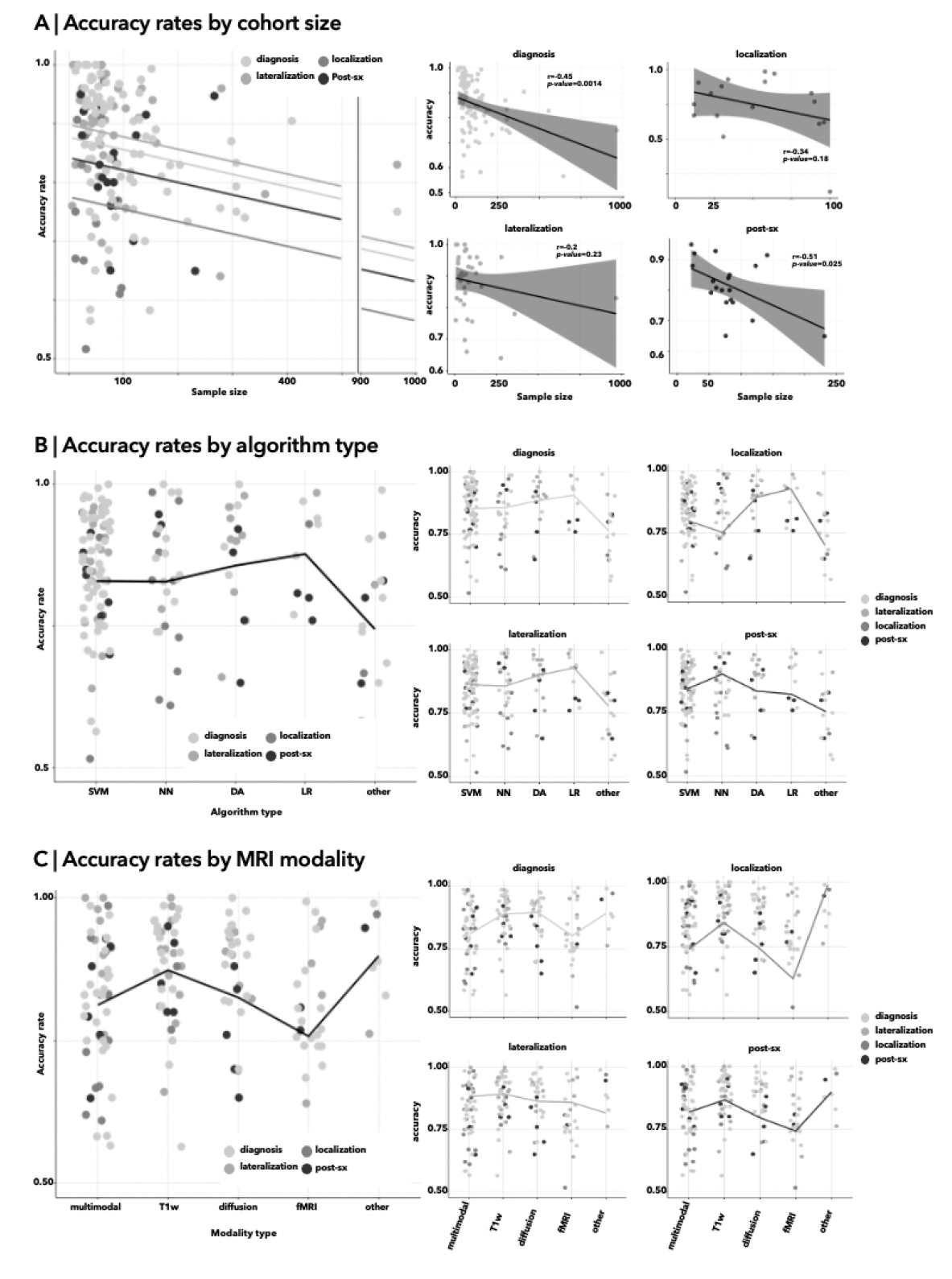
Comparing accuracy rates against study characteristics across outcomes. All models were fitte assuming mixed effects from the four main outcomes (diagnosis, lateralization, localization, and post-surgical outcome). (A) We assessed accuracy rate as a function of sample size, showing a negative association for all four clinical applications. (B) We evaluated accuracy rate by algorithm type, from most common to least common algorithm type. (C) MRI modality was divided into multimodal and the four most common unimodal scans; fMRI showed the lowest performance overall and performed most poorly for lesion localization of the four applications. Multimodal scans were shown to perform relatively similarly to unimodal sequences.

Among four most common algorithms (SVM, NN, DA, LR) across outcomes, the highest reported accuracy rates were associated with a LR-based model, followed by NN and SVM (**Figure 3B**). When examining individual outcomes, LR maintained increased accuracy for the purposes of diagnosis, localization, and lateralization compared to other algorithms. While SVM and NN performed relatively similarly across most outcomes, NN underperformed compared to SVM in being able to localize lesions despite its most common usage for this application.

We also evaluated whether the most common MRI sequences (five categories: multimodal, T1-weighted [T1w], diffusion, functional MRI [fMRI], and ‘other’ [FLAIR, qT1]) affected accuracy rates (**Figure 3C**). Notably, accuracy rates for publications using multimodal MRI data did not exceed those using any single MRI modality, though comparisons were performed between studies and were greatly limited by heterogeneity in cohorts, imaging parameters and combinations, and methodology.

Finally, regression analyses of all stated study characteristics (cohort size, all algorithm types, unimodal versus multimodal MRI sequence) indicated that larger cohort sizes were highly associated with lower accuracy rates. LR and DA models demonstrated the strongest model performance, with little difference between studies using unimodal versus multimodal MRI.

### Risk of bias

All papers had a high risk of bias overall and across each of PROBAST’s four domains (participants, predictors, analysis, and outcome) (**Figure 4**). Participant bias was reported at 87% (67/76) for diagnostic studies, 64% (24/37) for lateralization studies, 88% (32/36) for localization studies, and 69% (18/26) for post-surgical outcome studies.

**Figure 4.**
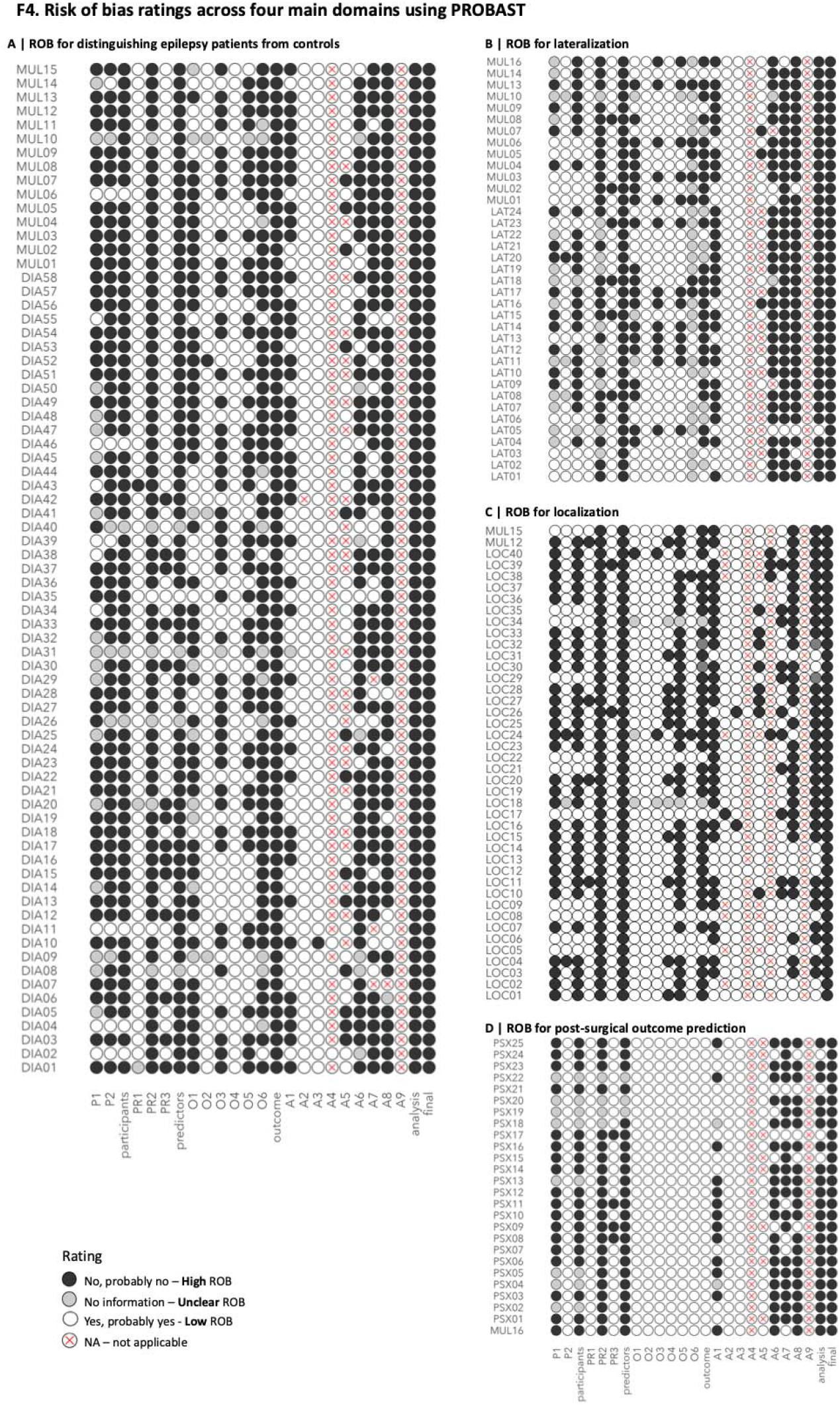
Comparing risk of bias (ROB) ratings using PROBAST across the four main clinical applications. All study IDs are listed on the left side of the matrix. The colour gradient of the matrix circles correspond to the risk of bias ratings; specifically, the darker the colour, the higher the risk of bias; circles with an X indicate that the question is not applicable (e.g. the question asks whether weights correspond to multivariate analysis results, but the study did not perform multivariate analysis). ROB assessment for each domain is determined by whether all domain questions are low risk of bias, and the overall ROB judgement is determined by whether all domains are considered low risk of bias. Hence, all studies were deemed to have an overall high risk of bias across all four outcomes.

High participant bias was attributed to the lack of appropriate recruitment; patients should be recruited prospectively at a time when the outcome is still being determined as opposed to retrospectively using data after definitive clinical diagnosis. For example, if the purpose is to distinguish epilepsy patients from controls, MRI data should be acquired during the diagnostic investigation as opposed to after definitive diagnosis. To be clinically useful, studies would ideally involve differential diagnostic tasks (such as the differentiation of people with epileptic seizures from those with functional seizures only), but study design may be challenged by sample size limitations, cohort heterogeneity, and the presence of co-occurring psychiatric comorbidities. Likewise, appropriate participant recruitment should occur prospectively for post-surgical seizure freedom prediction, as opposed to retrospectively, to prevent selection bias (*e*.*g*. patients who are seizure-free may be more likely to be lost to follow-up). Potential for biased predictors and analyses were similarly high across all outcomes since assessments were mainly made retrospectively and models were trained on small cohorts, here heuristically defined as less than 100 total participants (and a minimum of 50 patients, 50 controls) with no independent test cohort to validate performance metrics. Indeed, all localization and lateralization studies had predictor bias, while 88% (23/26) of studies predicting post-surgical outcome and 89% (67/76) of studies evaluating diagnostic value had high predictor bias. Similarly, all diagnostic studies had high analysis bias, and high analysis bias was also present in 97% (36/37) of lateralization studies, 83% (30/36) of localization studies, and 84% (22/26) of surgical outcome studies. Some questions were not applicable as most studies did not have missing data and therefore did not need to adjust for this (column A4) or did not use multivariate analysis to assign weights to predictors (column A9) and were therefore excluded from the overall judgement for risk of analysis bias. The risk of bias for the outcomes domain was low in post-surgical prediction studies, as seizure freedom is a clearly defined outcome, but remained high for diagnosis (100% [73/73]), localization (86% [31/36]), and lateralization (97% [36/37]) applications due to the use of overlapping MRI findings both as part of the clinical evaluation and as input to AI/ML algorithms, particularly for studies that used T1w MRI scans.

## DISCUSSION

With the advent of AI and ML methods, there has been increasing interest towards applying these techniques to multimodal MRI to effectively support epilepsy diagnosis and management. Our systematic review and meta-analysis synthesized this rapidly growing literature across four key clinical applications and highlights key avenues as well as challenges to clinical translation. A total of 158 papers with 26,732 participants (n=9,928 patients) were aggregated, underscoring a rapidly growing field of clinically oriented epilepsy research. The accuracy of ML models was high across the outcomes under study, namely epilepsy diagnosis (8%), TLE lateralization (90%), lesion localization (82%), and post-surgical outcome prediction (83%). Linear mixed effects models showed that algorithm type and cohort size were significantly related to accuracy rate. While these results are promising, subsequent systematic risk of bias assessment based on the PROBAST framework also highlighted several noteworthy limitations in study design, conduct, and analysis across the existing literature, which may pose barriers to robust and practical translation.

Among study design choices, sample size was shown to be the most significant factor associated with reduced model accuracy. High accuracy in small samples may be reflective of potential overfitting, particularly when overly optimistic classifier evaluation schemes are used. While small sample settings thus relate to risk of bias, these findings also indicate a prevailing issue regarding systemic barriers to patient recruitment. Indeed, patients often wait years, if not decades prior to referral to tertiary care centers for surgical evaluation and intervention, whereby they may also participate in research studies with generally more optimized, high-quality MRI sequences.^6,8^ This inadvertently reduced selection of patients may fail to account for effects from key biological factors such as age and sex or to distinguish between effects from disease duration, medication, and pre-existing epilepsy-related impairment, ultimately limiting model generalizability and lowering confidence in performance. In addition to standardizing MRI investigations and to bringing these to earlier disease stages as previously recommended, we propose that clinical pipelines should be streamlined to improve the referral pipeline from primary to tertiary care centers, thus reducing wait times for patients and helping research studies at those centers become more generalizable and representative of the population of people with epilepsy, leading to greater improvements in future care.^10^

Indeed, the effects of cohort size may have eclipsed potential influences of classifier choice and MRI modality. Although our analyses indicate that classifier choice and MRI multimodality did not markedly influence accuracy, there is notable work and clinical consensus that suggest otherwise. There is now established evidence that the combined use of T2-FLAIR and T1-weighted images greatly improves TLE lateralization and FCD localization.^22–24^ The lack of findings in the current assessment may be due to cross-study heterogeneity that cannot be fully accounted for by the meta-analysis techniques we employed here. Future work to systematically compare classifier or MRI sequence(s) combinations on accuracy rates within and across studies will be needed to more sensitively reveal the expected advantage of multimodal MRI compared to univariate analysis. Overall, our results indicate that the field must progress beyond continued recombination of MRI modalities, preprocessing techniques, and algorithm types to increase model accuracy. More consideration must be placed, firstly, on ensuring sufficiently large cohort sizes, and secondly, on designing studies taking account of generalizability and clinical feasibility while maintaining strong model performances.

To improve generalizability of future studies, our risk of bias evaluation has identified several sources to address. Firstly, the timing of patient recruitment and data acquisition would ideally be prospective. Most retrospective datasets include the same MRI data used for diagnosis or lesion identification. However, using the same data to define the ground truth (*e*.*g*. left vs. right TLE, epileptogenic lesion, *etc*.) as a predictor for models could lead to inflated accuracy rates, particularly given that MRI findings are, and should be, integral to epilepsy diagnosis.^10,25^ Several independent reference standards in addition to MRI can be used, such as including only patients with long-term Engel IA outcomes after surgery and/or histological verification.. However, application of these criteria may reduce sample size and lead to a reduced representation of the patient population to those patients that undergo successful resective epilepsy surgery. Furthermore, despite their validity, these standards present their own set of challenges, as rates of seizure freedom tend to decrease over time, and histopathological results have been shown to vary across sections.^9,26^ Though perhaps no perfect “ground truth” exists, a multi-pronged approach to define it will certainly strengthen the reliability of reported results. Secondly, most patients recruited at tertiary care centers have decades-long disease durations. As cortical and subcortical changes may likely progress over the disease duration, long durations may bias models to identify features most affected long-term as opposed to features that may be most discriminatory upon earlier evaluation.^23–25^ Thirdly, the majority of studies were at risk for overestimation of model performance, as they did not account for key biological and socio-demographic covariates such as age and sex, used leave-one-out cross-validation, and did not comprehensively report model performance metrics. These sources of bias can be addressed concomitantly with expanding cohort size. Stated previously, the clinical referral pipeline can be streamlined by encouraging early referral to tertiary care centers after pharmaco-resistance is confirmed and by reducing socio-environmental barriers to specialized care.^29^ Not only would MRI data from patients referred earlier into their diagnostic journey be ideal for further diagnostic classification (*i*.*e*. FCD identification, TLE lateralization) and surgical outcome prediction, cohort sizes may also increase enough to incorporate greater patient heterogeneity and allow for normative modeling of biological covariates.^27– 29^ Finally, studies should also comprehensively report all performance metrics (*i.e*., sensitivity/specificity, precision/recall, overall accuracy, and AUC under ROC) or raw confusion matrices for future systematic comparisons between studies demonstrating the utility of their AI/ML-driven models.

Certain promising models designed for both lesion detection in focal epilepsy and TLE lateralization have built towards clinical feasibility and interpretability for physician end-users.^15,16,26^ These studies included large multi-site cohorts, rigorously tested for model performance, and have considered leveraging data derived from standard MRI protocols, yet remain underutilized. Highly specific requirements for MRI sequences, complex data processing pipelines, and imperfect harmonization of data across centers may limit ease of applicability for widespread adoption. Thus, future efforts should not only focus on continuing prospective model validation, but also on collaborating closely with clinical and industry partners to identify optimal strategies for integration. Our results also encourage physicians to adhere to the ILAE-recommended and standardized MRI protocols and research groups to leverage these clinically acquired scans for external, prospective validation.^10^

Clinical applications of AI/ML in epilepsy have already demonstrated significant advancement, and future work to address the remaining sources of bias will help clinically integrate models to directly support patients. Given that current clinical guidelines do not solely use MRI for decision-making but interpret findings holistically with other diagnostic tools (*e.g*. semiology, EEG, PET), the integration of AI/ML models applied to MRI with these other modalities may also serve as a future research direction for this field. Nevertheless, future studies must prioritize robust study design and move towards clinical integration by emphasizing ease of clinical integration, interpretability, and external site validation. We encourage closer collaborations within the healthcare system and between care centers and research groups for wider patient recruitment, and we emphasize the need for inclusive, large, multi-center datasets that reflect real-world clinical diversity.

## Supporting information

Supplemental eReference file

