## Supplemental Tables for "The Use of Artificial Intelligence In Magnetic Resonance Imaging of Epilepsy: A Systematic Review and Meta-Analysis"

Fig S1.

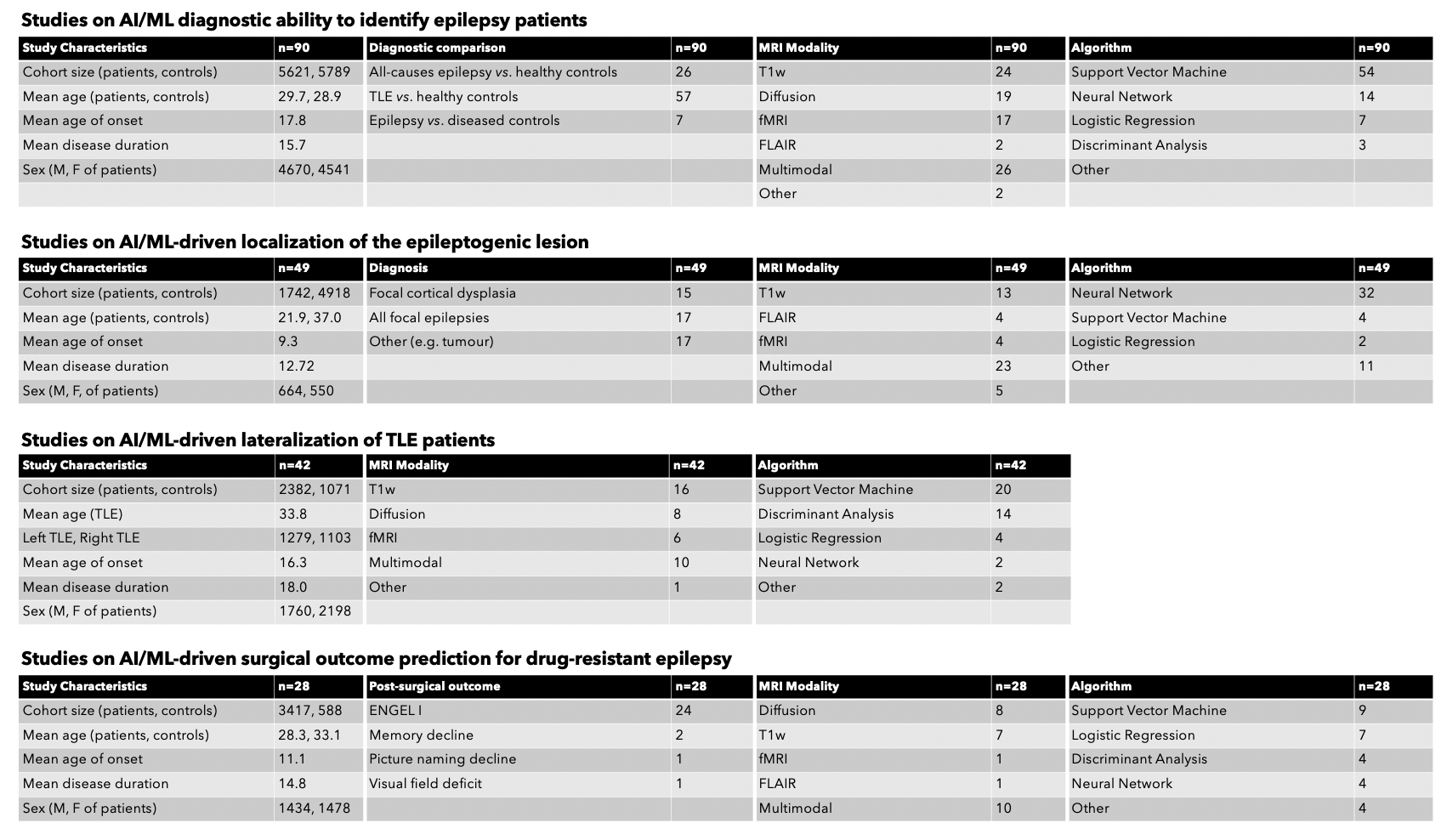

Table 1.1: Study IDs on AI/ML diagnostic ability and their corresponding citation (author, year).

| Hofer_2020 | DIA01 |
| --- | --- |
| Kerr_2014 | DIA02 |
| Zhou_2020 | DIA03 |
| Moguilner_2021 | DIA04 |
| Mueller_2013 | DIA05 |
| Shi_2021 | DIA06 |
| Garcia-Ramos_2023 | DIA07 |
| Chu_2022 | DIA08 |
| Huang_2020 | DIA09 |
| Casella_2023 | DIA10 |
| Nguyen_2021 | DIA11 |
| Nguyen_2020 | DIA12 |
| Huang_2022 | DIA13 |
| Zhu_2021 | DIA14 |
| Wu_2022 | DIA15 |
| Xie_2023 | DIA16 |
| Gonzalez_2021 | DIA17 |
| Tang_2022 | DIA18 |
| Rajpoot_2015 | DIA19 |
| Huang_2020 | DIA20 |
| Munsell_2015 | DIA21 |
| Mazrooyisebdani_2020 | DIA22 |
| Rudie_2015 | DIA23 |
| FerNIndez_2021 | DIA24 |
| Lee_2021 | DIA25 |
| Gaizo_2017 | DIA26 |
| Lee_2021 | DIA27 |
| Jiang_2023 | DIA28 |
| Park_2018 | DIA29 |
| Zhu_2022 | DIA30 |
| Chen_2020 | DIA31 |
| Park_2020 | DIA32 |
| Hwang_2019 | DIA33 |
| Kamiya_2016 | DIA34 |
| Gleichgerrcht_2021_brain | DIA35 |
| Hao_2022 | DIA36 |
| Mo_2021 | DIA37 |
| Yin_2023 | DIA38 |
| Ito_2021 | DIA39 |
| Cantor-Rivera_2014 | DIA40 |
| Chang_2023 | DIA41 |
| Jhuang_2020 | DIA42 |
| Si_2021 | DIA43 |
| Hogan_2006 | DIA44 |
| Hong_2017 | DIA45 |
| Zhang_2023 | DIA46 |
| Winston_2017 | DIA47 |
| Wang_2019 | DIA48 |
| Hao_2022 | DIA49 |
| Lai_2017 | DIA50 |
| Kang_2021 | DIA51 |
| Azzony_2023 | DIA52 |
| Su_2024 | DIA53 |
| Bharath_2019 | DIA54 |
| Amarreh_2014 | DIA55 |
| Gao_2022 | DIA56 |
| Guo_2020 | DIA57 |
| Gupta_2014 | DIA58 |
| Rebsamen_2022 | DIA59 |
| Raj_2010 | DIA60 |
| Gleichgerrcht_2021 | MUL01 |
| Gholipour_2021 | MUL02 |
| McDonald_2008 | MUL03 |
| Fang_2015 | MUL04 |
| Fang_2017 | MUL05 |
| Princich_2021 | MUL06 |
| An_2014 | MUL07 |
| Behesti_2021 | MUL08 |
| Focke_2012 | MUL09 |
| Kodipaka_2007 | MUL10 |
| Hong_2016 | MUL11 |
| Ganji_2022 | MUL12 |
| Sarbisheh_2022 | MUL13 |
| Johnson_2022 | MUL14 |
| Kulaseharan_2019 | MUL15 |

Table 1.2 Study IDs on AI/ML-driven lateralization of TLE patients and their corresponding citation (author, year).

| Pustina_2015 | LAT01 |
| --- | --- |
| Bennett_2019 | LAT02 |
| Caldairou_2021 | LAT03 |
| Beheshti_2020 | LAT04 |
| Kaestner_2023 | LAT05 |
| Bernhardt_2011 | LAT06 |
| Jamali-Dinan_2020 | LAT07 |
| Yang_2015 | LAT08 |
| Kamiya_2015 | LAT09 |
| Mo_2019 | LAT10 |
| Fallahi_2020 | LAT11 |
| Ahmadi_2009 | LAT12 |
| Farid_2012 | LAT13 |
| Keihaninejad_2012 | LAT14 |
| Fu_2021 | LAT15 |
| Duchesne_2006 | LAT16 |
| Lai_2017 | LAT17 |
| Xie_2022 | LAT18 |
| Chiang_2014 | LAT19 |
| Jafari-Khouzani_2009 | LAT20 |
| Mahmoudi_2018 | LAT21 |
| Barron_2014 | LAT22 |
| Wang_2023 | LAT23 |
| García-Pallero_2019 | LAT24 |
| Gleichgerrcht_2021 | MUL01 |
| Gholipour_2022 | MUL02 |
| McDonald_2008 | MUL03 |
| Fang_2015 | MUL04 |
| Fang_2017 | MUL05 |
| Princich_2021 | MUL06 |
| An_2014 | MUL07 |
| Beheshti_2021 | MUL08 |
| Focke_2012 | MUL09 |
| Kodipaka_2007 | MUL10 |
| Sarbisheh_2022 | MUL13 |
| Johnson_2022 | MUL14 |
| Hong_2016 | MUL16 |

Table 1.3 Study IDs on AI/ML-driven lesion localization and their corresponding citation (author, year).

| House_2021 | LOC01 |
| --- | --- |
| Hunyadi_2015 | LOC02 |
| Alaverdyan_2020 | LOC03 |
| Kanber_2021 | LOC04 |
| Jeong_2022 | LOC05 |
| Adler_2016 | LOC06 |
| ElAzami_2016 | LOC07 |
| Gill_2021 | LOC08 |
| Mo_2022 | LOC09 |
| Wang_2019 | LOC10 |
| Boerwinkle_2022 | LOC11 |
| OliveiraBaffa_2022 | LOC12 |
| Ganji_2021 | LOC13 |
| Gyebnár_2019 | LOC14 |
| Hong_2014 | LOC15 |
| Demerath_2021 | LOC16 |
| Sakashita_2023 | LOC17 |
| Marecek_2021 | LOC18 |
| BijayDev_2019 | LOC19 |
| David_2021 | LOC20 |
| Jin_2018 | LOC21 |
| Lin_2021 | LOC22 |
| Niyas_2021 | LOC23 |
| Qu_2016 | LOC24 |
| Spitzer_2022 | LOC25 |
| Snyder_2021 | LOC26 |
| Tan_2018 | LOC27 |
| Thomas_2021 | LOC28 |
| Wang_2018 | LOC29 |
| Urbach_2021 | LOC30 |
| Makhalova_2022 | LOC31 |
| Luckett_2022 | LOC32 |
| Nandakumar_2023 | LOC33 |
| Ganji_2022 | MUL12 |
| Kulaseharan_2019 | MUL15 |

Table 1.4 Study IDs on AI/ML-driven surgical outcome prediction for drug-resistant epilepsy and their corresponding citation (author, year).

| Benjumeda_2020 | PSX01 |
| --- | --- |
| Stasenko_2023 | PSX02 |
| Munsell_2015 | PSX03 |
| Larivière_2020 | PSX04 |
| He_2017 | PSX05 |
| Taylor_2018 | PSX06 |
| Pinheiro-Martins_2011 | PSX07 |
| Sinclair_2021 | PSX08 |
| Wang_2022 | PSX09 |
| Hinds_2023 | PSX10 |
| Sinha_2021 | PSX11 |
| Ko_2021 | PSX12 |
| Lee_2022 | PSX13 |
| Tang_2022 | PSX14 |
| Bernhardt_2014 | PSX15 |
| David_2021 | PSX16 |
| Wang_2023 | PSX17 |
| Memarian_2015 | PSX18 |
| Wagstyl_2021 | PSX19 |
| Johnson_2022 | PSX20 |
| Gleichgerrcht_2020 | PSX21 |
| Gleichgerrcht_2018 | PSX22 |
| Uijl_2008 | PSX23 |
| Yossofzai_2022 | PSX24 |
| Binding_2023 | PSX25 |
| Hong_2016 | MUL16 |
